# Uncovering Robust Patterns of MicroRNA Co-Expression across Cancers Using Bayesian Relevance Networks

**DOI:** 10.1101/115865

**Authors:** Parameswaran Ramachandran, Daniel Sánchez-Taltavull, Theodore J. Perkins

## Abstract

Co-expression networks have long been used as a tool for investigating the molecular circuitry governing biological systems. However, most algorithms for constructing co-expression networks were developed in the microarray era, before high-throughput sequencing—with its unique statistical properties—became the norm for expression measurement. Here we develop Bayesian Relevance Networks, an algorithm that uses Bayesian reasoning about expression levels to account for the differing levels of uncertainty in expression measurements between highly- and lowly-expressed entities, and between samples with different sequencing depths. It combines data from groups of samples (e.g., replicates) to estimate group expression levels and confidence ranges. It then computes uncertainty-moderated estimates of cross-group correlations between entities, and uses permutation testing to assess their statistical significance. Using large scale miRNA data from The Cancer Genome Atlas, we show that our Bayesian update of the classical Relevance Networks algorithm provides improved reproducibility in co-expression estimates and lower false discovery rates in the resulting co-expression networks. Software is available at www.perkinslab.ca/Software.html.

## Introduction

Co-expression of genes, microRNAs, long non-coding RNAs and other transcribed entities is a key biological property with multiple implications [1–4]. It can help identify functions of uncharacterized genes based on expression similarity to characterized genes [1]. Co-expression sometimes indicates co-regulation at the transcriptional level, thereby revealing how gene expression is controlled [2,5, 6]. Or, co-expression can be the result of coordinated epigenetic mechanisms [7, 8]. In yet other instances, co-expression of certain genes can serve as biomarkers in diseases such as cancer [9, 10] and mental disorders [8], or aid in defining distinct cell populations and subpopulations [11].

One of the earliest, and still widely used, tools for estimating and exploring networks of co-expression is the Relevance Networks algorithm of Butte *et al*. [12]. The algorithm has four main steps. First, entities (e.g., genes) with low estimated entropy are removed, as correlations between them may result from one or a few outlier samples. Second, Pearson correlations are computed between all pairs of remaining entities. Third, permutation testing is used to establish a null distribution for the correlations. Fourth and finally, a co-expression network is created by connecting any pair of entities whose correlation exceeds a statistical significance threshold set by the user (in conjuction with the estimated null distribution). The Relevance Networks algorithm has been used successfully in numerous studies to uncover significant co-expression relationships (e.g., [13–16]).

Many elaborations and alternatives to the original Relevance Networks algorithm have been proposed over the years [17–23]. These include improvements aimed at detecting non-linear relationships between the expression of different entities by using mutual information criteria, as seen in Mutual Information Relevance Networks and the ARACNE algorithm [17, 18, 24], or discriminating co-expression more likely to result from direct rather than indirect interactions, as seen in ARACNE and CLR [18, 20, 24]. As replicate data became more common, algorithms were developed to accomodate co-expression analysis of data with replicates, as in Zhu *et al*. and Acharya and Zhu [21, 23]. Other work has focussed on robustly estimating correlations when the number of samples is much smaller than the number of entities, as in Sch afer and Strimmer [19]—although interestingly, we are finally emerging from that conundrum. For instance, the dataset we analyze in this paper describes 2,456 miRNAs measured over 10,999 samples. Other tools, such as WGCNA, go beyond the construction of correlation networks, offering features of module identification, topology analysis, etc. [22].

While these algorithms embody many important ideas and methods for co-expression network construction and analysis, they were all developed in the era of microarray-based expression measurements. The recent past has seen a fundamental shift in the technology used for expression measurement from microarray-based to sequencing-based platforms [25, 26]. Sequencing-based approaches produce measurement values with very different error properties, dynamic ranges, and signal-to-noise ratios than microarrays. In particular, the relative precisions of low-expression measurements are much worse compared to those of high-expression measurements. Furthermore, precision differs between samples, at the very least due to differences in sequencing depth, if not other factors [27]. Intuitively, these differences in precision should influence our confidence in co-expression estimates. Indeed, in the context of microarrays, the pioneering work of Hughes et *al*. [1] clearly established the value of accounting for gene-specific measurement uncertainties in assessing co-expression and differential expression. How can the same concept be translated into the statistically much different setting of high-throughput sequencing data?

Here, we develop a Bayesian version of the classical algorithm of Butte et *al*. [12], which we call the Bayesian Relevance Networks algorithm. It builds on our recent work where we proposed a Bayesian correlation scheme to analyze sequence count data [28]. We employ Bayesian statistics both for estimating the expression levels and for quantifying the uncertainties in those estimates. From those beliefs, we construct estimates of mean expression levels and their uncertainties in groups of samples. This allows us to study cross-group correlations in studies with replicates or other natural sample groups (e.g., patients with the same disease). We describe how to perform permutation testing to estimate a null distribution for grouped Bayesian correlations. This enables the computation of *p*-values for the statistical significance of observed correlations, and allows us to estimate rates of true and false positive links in a Bayesian Relevance Network.

Throughout the paper, we evaluate our approach on a large-scale public microRNA (miRNA) expression dataset from The Cancer Genome Atlas project (TCGA) [29]. In a series of cross-validation studies, we find that Bayesian co-expression estimates are more reproducible than the Pearson co-expression estimates used by the original Relevance Networks algorithm. We find that Bayesian Relevance Networks are less prone to false positive links and have lower false discovery rates than classical Relevance Networks. Finally, we find that entropy filtering to remove “spurious” correlations improves both classical and Bayesian Relevance Networks. At the end of the Results section, we present a Bayesian Relevance Network based on the full datasets, where we demonstrate several interesting cancer type-specific clusters of co-expressed miRNAs.

## Materials and Methods

### Problem Formulation

The algorithm we propose is for computing a co-expression network among *m* possible entities (genes, miRNAs, etc.) measured across a set of samples organized into *n* groups. The groups may represent replicates of a condition, patients with a common disease, etc. Group *g* has *n_g_* samples in it.

We observe *R_igs_* reads for entity *i* in group *g* sample *s*. The total number of reads for that sample is 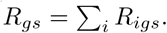. We assume that the *R_igs_, i* = 1… *m*, are multinomially distributed.

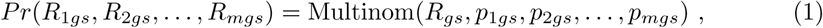
where the *p_igs_* are unknown. Each *p_igs_* represents the idealized fraction of the sample *s* in group *g* that comes from entity *i*. We can also think of it as what *R_igs_/R_gs_* should converge to in the limit of infinite sequencing depth (*R*_gs_ → ∞). We define the group mean idealized fractions as 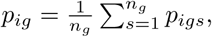, and the grand mean idealized fraction as 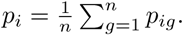.

We take the *p_igs_* to be our definition of the expression level. Other common definitions include reads per million (RPM), or fragments per kilobase per million (FPKM). Both of these normalize for sequencing depth in a given sample and are proportional to *p_igs_*. As correlations are independent of scale, working with the *p_igs_* is equivalent to working with RPM or FPKM. Other normalization schemes could be accomodated, as long as the expression level can be written as an affine function of the *p_igs_*. However, so as not to overly complicate our notation, we leave this to the reader.

For any entity *i*, we define the cross-group variance in expression as

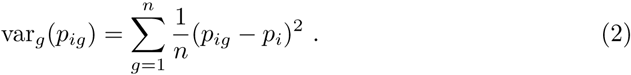

In this formula, we are essentially treating the group *g* as if it were a random variable, taking values 1… *n* with equal probability. For any two entities *i* and *j* we define the cross-group covariance as

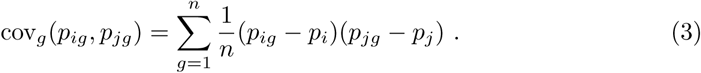

Then, the cross-group Pearson correlation of their expression values of entities *i* and *j* is defined as

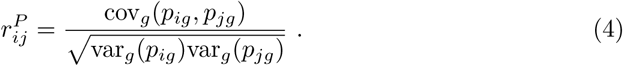

Ideally, we would like to connect entities *i* and *j* in a co-expression network if their cross-group correlation is statistically significantly large. The problem, of course, is that the *p_ig_* are unknown, so we must estimate them.

### The Bayesian Relevance Networks Algorithm

In principle, one could construct a Bayesian belief about the unknown Pearson correlation itself. However, this is not computationally convenient. Instead, we use Bayesian methods to construct estimates of the expression levels, *p_igs_*, and then estimate their correlations. The algorithm we propose has four steps, which are detailed in the following subsections.

1. Remove low entropy entities from consideration (optional).
2. Compute Bayesian estimates of the cross-group correlations of expression between every (remaining) pair of entities
3. Use permutation computations to estimate a null distribution for the Bayesian cross-group correlations
4. Create a network by linking entities whose Bayesian correlations are statistically significant

### Entropy filtering

This step is optional. We include it for the same reason it was included in the original Relevance Networks algorithm—that correlations may arise spuriously due to outliers. For instance, suppose two entities are generally expressed at constant levels, but in one sample both of their levels are much higher or lower than normal. These two entities will thus appear to have highly correlated expression levels. In some cases this may be genuinely true, although we may not be comfortable about the robustness of a correlation that depends on a single sample being present in the dataset. The same phenomenon might also arise for more mundane reasons, such as sample mishandling, contamination, poor sequencing depth, etc. Thus, it may make sense to remove entities with such expression profiles from consideration.

To allow for direct comparison between our Bayesian approach and the classic Relevance Networks algorithm, we use the exact same entropy filtering procedure. For each entity *i*, we compute the maximum likelihood expression estimates, 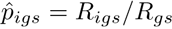. We then compute the minimum, 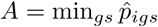, and maximum, 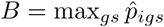, expression levels across all samples in all groups. If *A* = *B* then we estimate the entropy of entity *i*’s expression as *H_i_* = 0. Otherwise, we divide the interval [*A, B*] into 10 equal-sized bins. We determine the empirical fraction of the 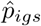 that fall into each of those 10 bins, calling them *f_i_*_1_…*f_i_*_10_. We then estimate the entropy of entity *i*’s expression as 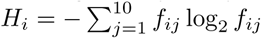 Entities with estimated entropies in the lowest *H_thresh_*% are discarded, where *H_thresh_* is chosen by the user.

### Bayesian estimation of pairwise correlations

The essence of our Bayesian approach is to first construct beliefs over the true expression levels of all the entities. We then propose that the Pearson correlation between two entities be replaced by what we call the Bayesian correlation. We compute variances and covariances across groups and also with respect to our uncertainty about the true expression levels. That uncertainty arises from the limited sampling depth in any experiment and the inherent noise in sampling reads from the large set of possible entities. Using *u* to denote our uncertainty informally—and we will become formal very shortly—the Bayesian correlation can be written as

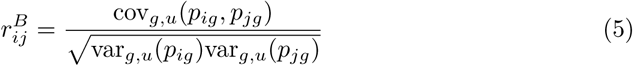

Intuitively, high uncertainty in expression levels may influence the covariance term, but it will definitely inflate the variance terms in the denominator, leading to lower estimates of correlation. (More precisely, estimates moderated towards zero.)

We adopt a standard Bayesian approach to estimate the idealized fractions *p_igs_*. For each group *g* and sample *s*, we employ a Dirichlet distribution to model our uncertainty about the *p_igs_*. We assume the Dirichlet beliefs for different samples are independent. Thus, for sample *s* and group *g* we adopt a prior belief,

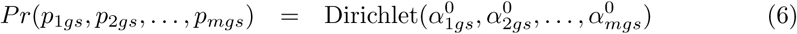

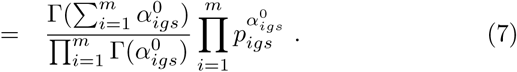

The posterior distribution is

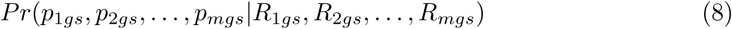

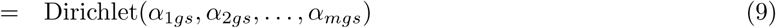

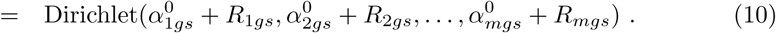

The prior parameters 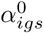 may be chosen however one likes. We previously showed that poor choice of priors can lead to highly biased estimates of correlation [28], and thus some care should be taken with the choice. We employ 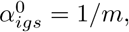, which has provably low bias for low expression entities represented by few read counts [28]. For entities with high read counts, the prior makes little difference, as the posterior is determined almost entirely by the data. With these assumptions, and defining 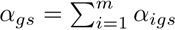, the mean of the marginal posterior distribution for *p_igs_* with respect to our beliefs (which we denote by *u* for “uncertainty”) is

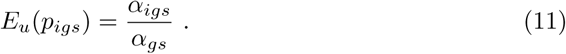

The variance of that marginal posterior is

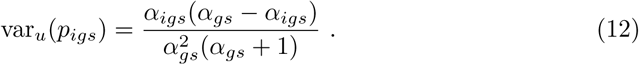

The covariance of our beliefs about the expression of two different entities, *i* and *j* ≠ *i*, within the same sample *s* of group *g* is

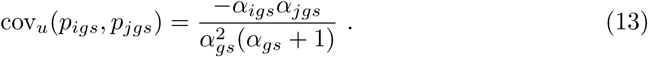

This covariance is nonzero because of the implicit requirement that 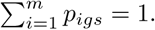. Intuitively, if we believe that *i*’s expression is larger, we must believe that the expression of other entities is smaller.

From these, we can readily compute the within-group means, variances and covariances between entities, accounting for our uncertainty. Recalling that by definition, *p_ig_* is the average of *p_igs_* across samples *s*, we have the following.

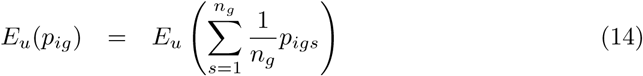

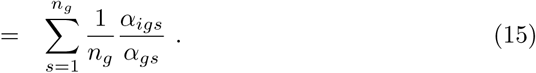

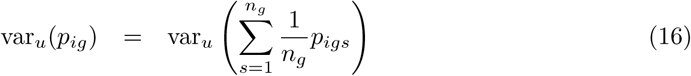

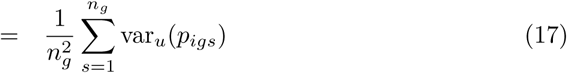

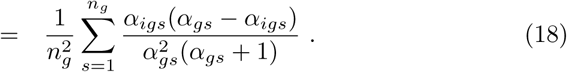

Eq 17 follows because our estimates for different samples are statistically independent, so the variance of the sum is the sum of the variances.

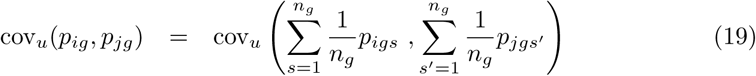

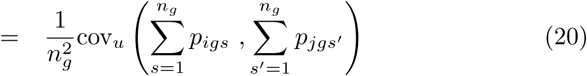

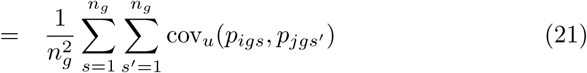

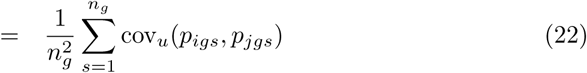

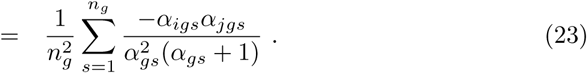

Eq 22 follows because our beliefs are independent for different samples, hence there is no covariance when *s* ≠ *s*′. We can then define the total variance across groups and uncertainty, for entity *i*, via the Law of Total Variance as

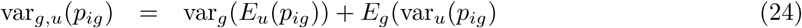

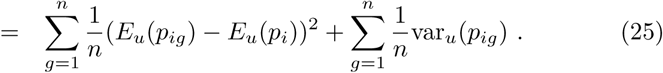

Similarly, we define the total covariance across groups and uncertainty, for entities *i* and *j*, via the Law of Total Covariance as

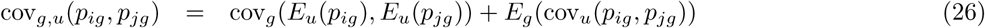

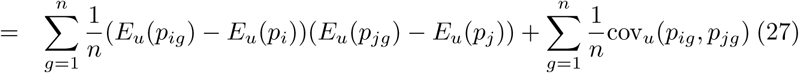

Eqs 25 and 27 can be substituted back into Eq 5 to completely specify the definition and computation of the Bayesian correlation. One step of this substitution and expansion is displayed below, as it will be relevant to our discussion of permutations in the next section.

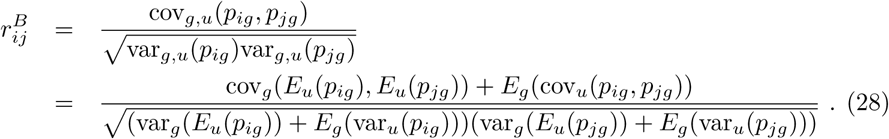

### A Permutation Scheme for Assessing Statistical Significance

Permutation testing is a common approach to assessing significance of associations between variables. However, in our context, this is not entirely straightforward. It is not sufficient to simply permute the read counts *R_igs_* for each entity *i* and recompute Bayesian correlations. Recall that the estimated expression levels of entity *i* depend not only on *R_igs_* but also on the total reads in the samples, *R_gs_*. Permuting the read counts would change the *R_gs_*, and therefore change the estimated expression levels. Permutation testing should “break” associations between different entities by reassigning their values to different samples, but it should not change the values themselves. It is also not sufficient to permute the estimated expression levels, *E_u_ p_igs_*, as that could change estimated group expression levels, *E_u_ p_ig_*.

With the null hypothesis being that there is no cross-group correlation between entities, we suggest that a proper way to estimate a null distribution between entities *i* and *j* is to compute many different permutations *ρ*: {1… *n*}↦{1… *n*} of the group numbers (all permutations, if possible). For each permutation *ρ* we evaluate the following formula.

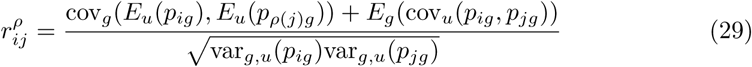

The distribution of that value for many different permutations *ρ* is taken to be the null distribution of the Bayesian correlation.

In comparison with the formula for the Bayesian correlation (Eq 28), the permuted values of *j*’s group-level expression are used in the first covariance term. This is the part of the formula where the hypothesis of no cross-group correlation would have its effect. We do not use the permuted *j*’s in the second covariance term. That term represents the covariance of our beliefs within a sample, which results from the necessity that expression levels within a sample add up to one. This is not affected by the null hypothesis, so we leave it unchanged. The permutations also do not appear in the variance terms of the denominator, although it would not matter if they did, as the variances of *i*’s and *j*’s expression are independent.

### Statistical Significance and Constructing the Bayesian Relevance Network

In the classical Relevance Networks algorithm, a single null distribution for correlations under the null hypothesis is constructed by combining the permuted correlations across all pairs of entities. Although it is technically more sound to maintain a separately estimated null distribution for each pair of entities (*i, j*), in order to maximize our ability to compare the results of Bayesian Relevance Networks to the classical algorithm, we do the same here. Thus, suppose that *K* times we have permuted the group idealized fractions, *E_u_ p_ig_*, of every entity *i*, and recomputed the cross-group Bayesian correlations as in Eq 29. Let 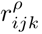 represent the permuted Bayesian correlation between entities *i* and *j* in the *k^th^* permutation. We estimate the overall probability of a correlation of at least *t*, under the null hypothesis, as

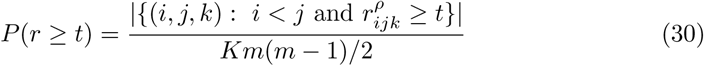

Suppose we construct a Bayesian Relevance Network by connecting any pair of entities *i* and *j* if their Bayesian correlation is at least *t*, obtaining *N_t_* such pairs. Given that there are *m*(*m* – 1)/2 possible pairs of entities, we can estimate the expected number of false positives at that threshold as *FP_t_* = *P*(*r* ≥ *t*)*m*(*m –* 1)/2. The number of true positives can be estimated as max(*N_t_* – *FP_t_*, 0). The false discovery rate can be estimated as min(*FP_t_/N_t_*, 1), as long as *N_t_* > 0. Together, these quantities—estimated numbers of true positives, numbers of false positives, and the false discovery rate—can be employed by the user to make a rational choice for the threshold *t* used to construct the network.

### Data

To demonstrate and evaluate our approach, and potentially to generate some biological insights in an important area, we decided to analyze miRNA expression data from The Cancer Genome Atlas (TCGA) [29]. We used the Genomic Data Commons data portal [30] to download all available “isoforms.quantification.txt” files on November 10, 2016. These files report counts of miRNA-seq reads mapped to a large number of genomic intervals. Those intervals are also annotated for whether they represent a certain pre-miRNA, a mature miRNA, or several other types of objects. From each file, we collected all lines corresponding to a mature miRNA (specified by a unique miRBase [31] MIMAT identifier), and then added up all counts corresponding to the same mature miRNA. This includes reads mapped to slightly different genomic intervals within the same mature miRNA, as well as entirely different genomic regions that happen to code for the same mature miRNA. In the end, this left us with read counts for 2456 distinct mature miRNAs, across 10,999 patient samples.

While this gave us a wealth of data on miRNA expression in cancer, the isoform files do not specify which types of cancer each patient had (nor any other patient characteristics). To establish this information, we constructed a json query that, through the Genomic Data Commons API, returned a list of all isoform quantification files, along with their project IDs. The project IDs are synonymous with the types of cancer profiled. In this way, we assigned one of 33 unique cancer types to each miRNA-seq dataset. These cancer types consititute the groups in our grouped correlation analysis.

In order to better inform our co-expression assessments, we downloaded from miRbase [31] their version 21 miR definitions in the file “hsa.gff3”. This file specifies the IDs and genomic coordinates of both stem-loop pre-cursors and mature miRNAs. It also specifies which mature miRNAs are to be found in which stem-loop precursors. Multiple genomic occurrences of the same mature miRNA have IDs ending in _1, _2, etc., to discriminate them. However, the “Alias” field omits these IDs, which could then be matched to the MIMAT IDs in the TCGA isoforms file. Similarly, we downloaded from ENSEMBL their latest gene definitions in the file “Homo_sapiens.GRCh38.86.gtf”. This file describes many types of transcribed entities, including protein-coding genes, pseudogenes, long non-coding RNAs, miRNAs, etc. Importantly, it includes their genomic locations. Using these sources of information, we were able to categorize every pair of mature miRNAs into one of the following categories: (1) “stem-loop” if the two mature miRNAs occur within the same stem-loop precursor miRNA anywhere in the genome; (2) “transcript” if the two mature miRNAs occur within the same transcribed entity (according to ENSEMBL) but not the same stem-loop precursor; (3) “near” if the two mature miRNAs occur within 10kb on the genome; (4) “cluster” if the two mature miRNAs occur within the same equivalence class in the transitive closure of the “near” relation, but are not themselves “near”. For example, if *i* is near *j* and *j* is near *k*, but *i* and *k* are not near, then *i* and *k* are still in the same cluster; (5) “non-local” if none of the previous categories apply.

## Results

### TCGA miRNA expression data spans many orders of magnitude across miRNAs and samples

As described in the Methods section, we obtained miRNA-seq expression data from the TCGA project through the Genomic Data Commons, resulting in read counts for 2456 miRNAs in 10,999 patient samples, representing 33 cancer types. Some summary statistics are shown in Table 1, while the data is shown visually in Fig 1A. Each row corresponds to a miRNA, and each column corresponds to a patient sample. The most-represented cancer was breast cancer, with 1207 samples, while the least-represented was glioblastoma multiforme, with 5 samples. There are clearly miRNAs with cancer-specific, or at least tissue-specific, expression profiles. To formalize this observation, we computed for each miRNA the total variance across samples of its expression in units of reads per million. We also computed the variance of expression within and across cancer types, and from those the percentage of variance explained (POVE) for each miRNA by the cancer type. Figure 1B shows a histogram of the POVE for all miRNAs. While for most miRNAs patient-to-patient variability within cancer types is dominant, there is a subset for which differences between cancer types are substantial. For approximately 3% of miRNAs, differences between cancer types explain the majority (≥ 50%) of the variability.

**Fig 1.**
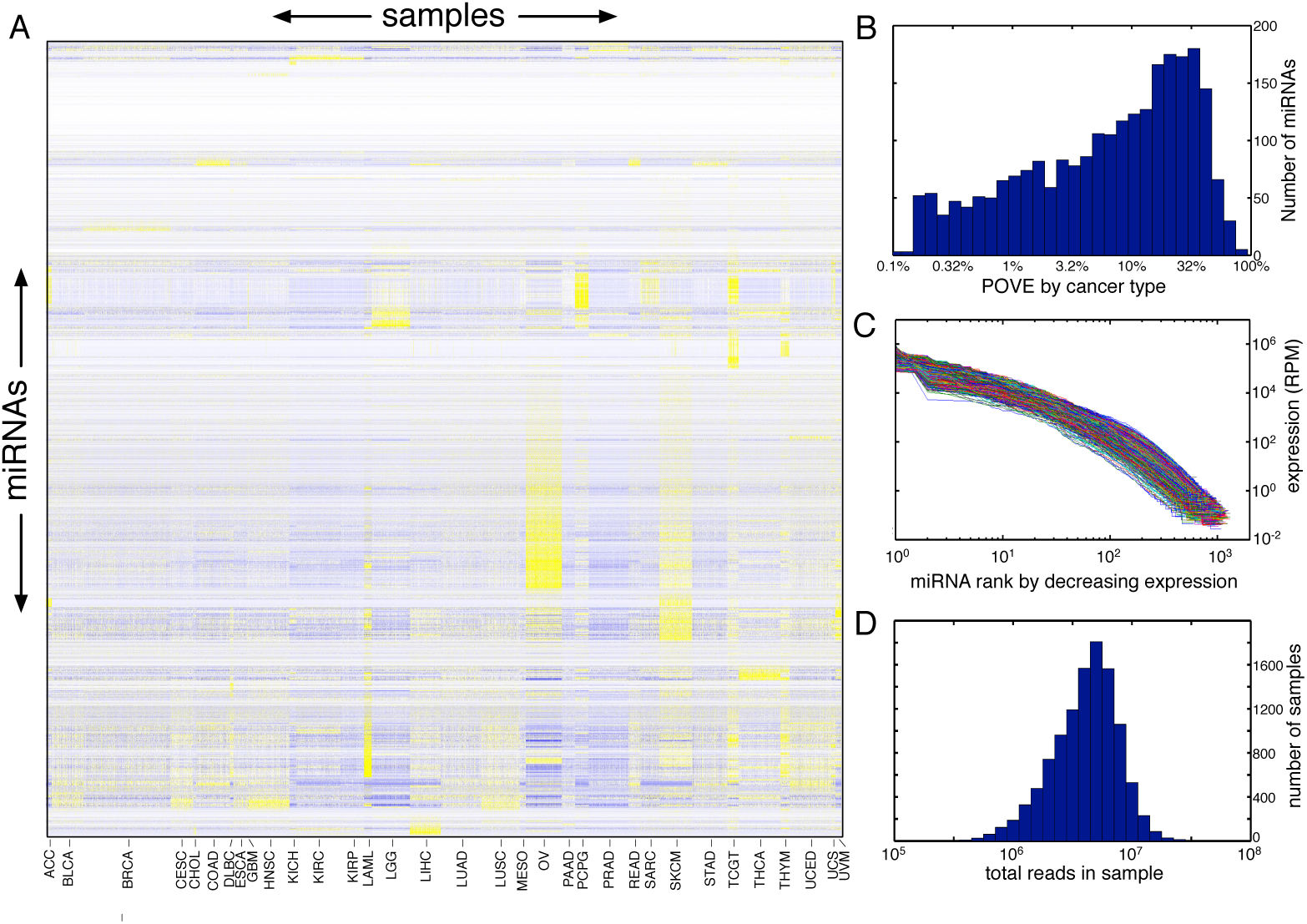
Mature miRNA expression data for 10,999 cancer patients from the TCGA project. (A) Heatmap of expression, with yellow indicating high and blue indicating low, relative to the mean for each miRNA across samples. miRNAs are ordered based on a hierarchical average-linkage Euclidean-distance clustering of the reads per million across samples. Samples are grouped by cancer type, indicated by labels along the bottom. (B) Histogram of percentage of variability in expression of different miRNAs explained by differences in cancer type. (C) Curves showing expression of all miRNAs within each sample, sorted from highest to lowest expression. (D) Histogram of the numbers of reads (i.e., sequencing depth) in each sample.

**Table 1.**
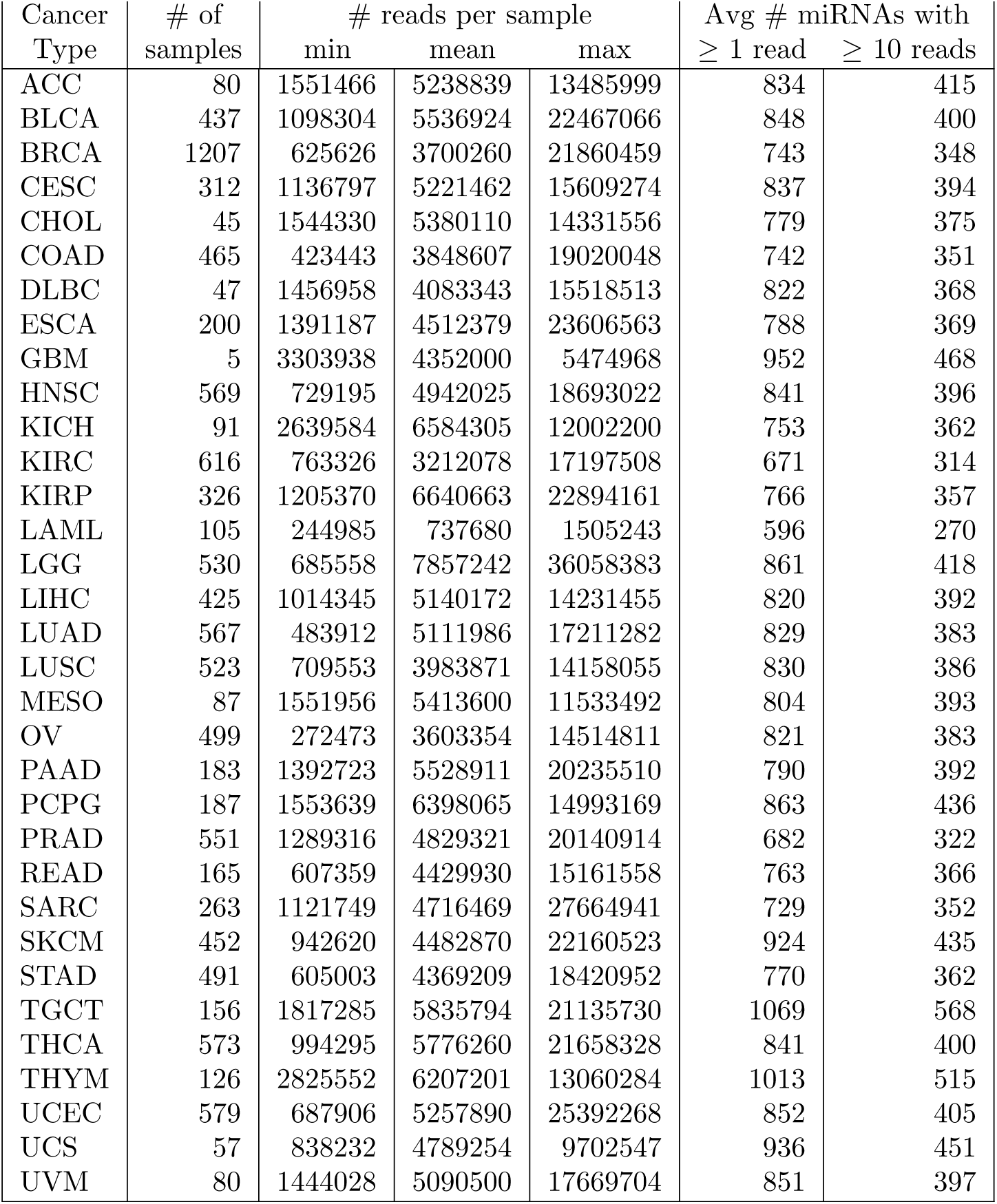
Summary statistics of the TCGA miRNA expression data, arranged by the 33 cancer types indicated in the first column.

| Cancer Type | # of samples | # reads per sample |  |  | Avg # miRNAs with |  |
| --- | --- | --- | --- | --- | --- | --- |
| | | min | mean | max | $\geq 1$ read | $\geq 10$ reads |
| ACC | 80 | 1551466 | 5238839 | 13485999 | 834 | 415 |
| BLCA | 437 | 1098304 | 5536924 | 22467066 | 848 | 400 |
| BRCA | 1207 | 625626 | 3700260 | 21860459 | 743 | 348 |
| CESC | 312 | 1136797 | 5221462 | 15609274 | 837 | 394 |
| CHOL | 45 | 1544330 | 5380110 | 14331556 | 779 | 375 |
| COAD | 465 | 423443 | 3848607 | 19020048 | 742 | 351 |
| DLBC | 47 | 1456958 | 4083343 | 15518513 | 822 | 368 |
| ESCA | 200 | 1391187 | 4512379 | 23606563 | 788 | 369 |
| GBM | 5 | 3303938 | 4352000 | 5474968 | 952 | 468 |
| HNSC | 569 | 729195 | 4942025 | 18693022 | 841 | 396 |
| KICH | 91 | 2639584 | 6584305 | 12002200 | 753 | 362 |
| KIRC | 616 | 763326 | 3212078 | 17197508 | 671 | 314 |
| KIRP | 326 | 1205370 | 6640663 | 22894161 | 766 | 357 |
| LAML | 105 | 244985 | 737680 | 1505243 | 596 | 270 |
| LGG | 530 | 685558 | 7857242 | 36058383 | 861 | 418 |
| LIHC | 425 | 1014345 | 5140172 | 14231455 | 820 | 392 |
| LUAD | 567 | 483912 | 5111986 | 17211282 | 829 | 383 |
| LUSC | 523 | 709553 | 3983871 | 14158055 | 830 | 386 |
| MESO | 87 | 1551956 | 5413600 | 11533492 | 804 | 393 |
| OV | 499 | 272473 | 3603354 | 14514811 | 821 | 383 |
| PAAD | 183 | 1392723 | 5528911 | 20235510 | 790 | 392 |
| PCPG | 187 | 1553639 | 6398065 | 14993169 | 863 | 436 |
| PRAD | 551 | 1289316 | 4829321 | 20140914 | 682 | 322 |
| READ | 165 | 607359 | 4429930 | 15161558 | 763 | 366 |
| SARC | 263 | 1121749 | 4716469 | 27664941 | 729 | 352 |
| SKCM | 452 | 942620 | 4482870 | 22160523 | 924 | 435 |
| STAD | 491 | 605003 | 4369209 | 18420952 | 770 | 362 |
| TGCT | 156 | 1817285 | 5835794 | 21135730 | 1069 | 568 |
| THCA | 573 | 994295 | 5776260 | 21658328 | 841 | 400 |
| THYM | 126 | 2825552 | 6207201 | 13060284 | 1013 | 515 |
| UCEC | 579 | 687906 | 5257890 | 25392268 | 852 | 405 |
| UCS | 57 | 838232 | 4789254 | 9702547 | 936 | 451 |
| UVM | 80 | 1444028 | 5090500 | 17669704 | 851 | 397 |

Fig 1C shows the expression of every miRNA in every sample, sorted by decreasing order within the sample. Expression values range from around 10^5^ RPM to below 1 RPM. Because all miRNAs are measured in the same units—reads—this means that relative to their expression levels, the miRNAs with lowest expression are measured with approximately 1/100,000 the precision of the miRNAs with highest expression. There are also great differences in sequencing depth between samples, as shown in Fig 1D. The sample with the greatest sequencing depth has over 36 million reads, while the sample with the shallowest sequencing depth has under a quarter million. There is approximately a 150-fold difference in resolution between these two samples. Given these statistics, it is clear that our uncertainties about the true expression levels of the miRNAs must vary widely by miRNA and by sample.

### Bayesian correlations are more reproducible than Pearson correlations

We expected that Bayesian correlation estimates would suppress correlations between low expression miRNAs. By contrast, we expected that Pearson correlations would be more subject to falsely high or low correlations, due to spurious correlations between miRNAs with low read counts. To test this, we computed all pairwise grouped Bayesian and Pearson correlations, using the cancer types to define patient groups. For the Pearson correlations, this was the correlation across cancers of the within-cancer average expression in units of RPM. Fig 2A shows a density scatterplot of the Pearson and Bayesian correlations. Points along the *y* = *x* diagonal line correspond to miRNA pairs where Pearson and Bayesian estimates agreed. We note that there are some miRNA pairs correlated at essentially +1 by both Pearson and Bayesian estimates, but no miRNA pairs with such strong anticorrelations. At the same time, there are many miRNA pairs that have high correlations according to the Pearson estimate, but that are relegated to much lower correlation levels—including essentially zero—by the Bayesian estimate. These involve miRNAs that, by our approach, have too much uncertainty in their expression levels to be able to confidently assert a strong correlation. As a rather extreme example, there was a strong disagreement in the estimated correlations between miR-4459 and miR-5692b. The former shows expression in 70 different samples across 12 cancer types, but is primarily seen in thyroid cancers, albeit at low levels (53 samples, 139 total reads). The latter is expressed at only 2 reads in a single thyroid cancer sample, and nowhere else. The Pearson correlation between these two is a near perfect 0.9731, whereas the Bayesian correlation is 0.0512.

**Fig 2.**
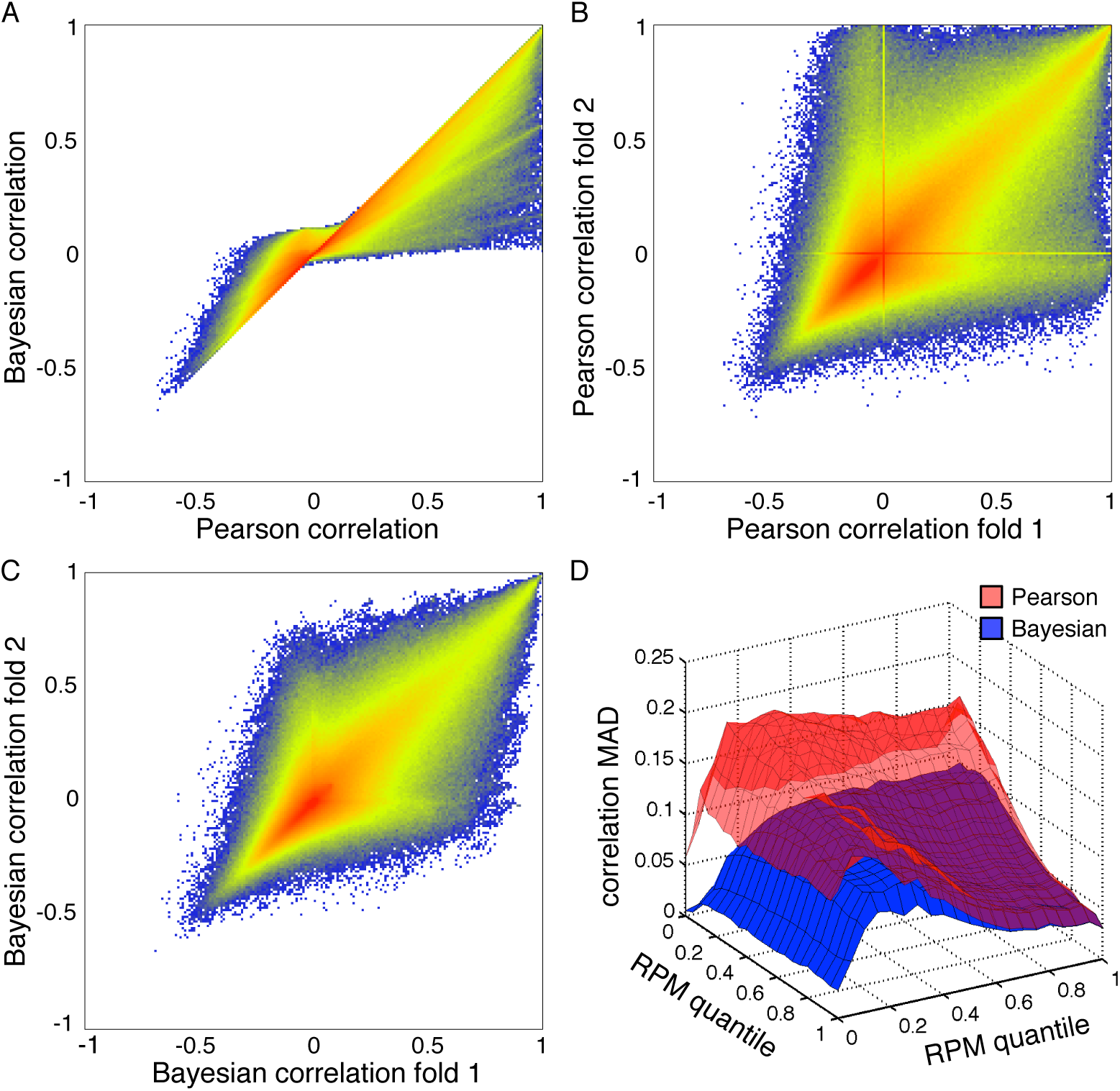
Comparison of Pearson and Bayesian grouped correlations across cancer types. (A) Density scatterplot of Bayesian versus Pearson correlations. Non-white points are where at least one pair of miRNAs has the specified Pearson (x-axis) and Bayesian (y-axis) correlations. Colored points, going from blue to yellow to red, indicate increasing numbers of miRNA pairs with the specified correlations. (B) Agreement of Pearson correlations when the data is divided in half and correlations computed for each half separately. (C) Agreement of Bayesian correlations when the data is divided in half and correlations computed for each half separately. (D) For each pair of miRNAs, organized by their expression quantiles across all samples, the average mean absolute deviation (MAD) between the two data halves of Pearson and Bayesian correlations.

To test the reproducibility of Pearson and Bayesian correlations, we randomly assigned each sample to one of two data folds, keeping the numbers of samples representing each cancer type as even as possible. We then computed cross-cancer Pearson correlations on each half of the data separately (Fig 2B), and likewise for the Bayesian correlations (Fig 2C). For the Pearson correlations, there is broad agreement between correlations computed based on each fold of the data—the estimates from each half are themselves correlated. But there are also many miRNA pairs where correlations from the two folds disagree dramatically. For a substantial number of pairs, one fold of the data produces a Pearson correlation near 1, while the other fold produces a Pearson correlation near zero. The two “lines” visible along the x- and y-axes of the density scatterplot arise from miRNAs that have absolutely zero reads in one fold of the data (hence no correlation to anything), but some reads in the other fold (and in some cases strong correlations, although they may be spurious). In comparison, the Bayesian correlation estimates from each fold of the data tend to be closer to each other. There are no “lines” of exceptional behaviour for zero-count miRNAs, and no miRNA pairs with near zero Bayesian correlation in one fold and near +1 Bayesian correlation in the other fold (although there are a very few near 0.9).

To quantify the reproducibility of the two approaches more carefully, and also to study the relationship between expression level and correlations, we divided miRNAs into 21 bins of increasing average RPM expression. Let *X* denote the set of miRNAs in one expression bin, and *Y* denote the set of miRNAs in another expression bin. From data fold 1, we computed all pairwise Pearson correlations between miRNAs in bin *X* with those miRNAs in bin *Y*, namely, 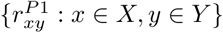. We did the same for data fold 2, compute the correlations 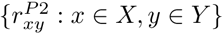. Finally, we computed the mean absolute deviation between these two sets of correlations, 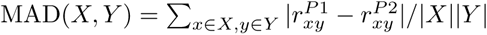 This gives the average disagreement of Pearson correlations computed from the two data folds, as a function of binned expression level. Then, we did the same for the Bayesian correlations. Fig 2D shows those mean absolute deviations. Generally, as the expression of both miRNAs trends higher, the disagreement between the two halves of the data decreases, and the error in the Pearson and Bayesian estimates is essentially identical. For these miRNAs, low signal-to-noise ratio is not an issue, and Pearson and Bayesian estimates are nearly the same. Error is worst when both miRNAs have low but nonzero expression, and it is nearly as bad when just one of the two miRNAs has low but nonzero expression. When one of the miRNAs is in the lowest expression bin, error tends not to be quite as bad, as both methods will tend to assign zero correlation (but Bayesian more so than Pearson). At all levels of expression, the average error of the Pearson estimates exceeds the error of the Bayesian estimates. Across all pairs of miRNAs, the Pearson MAD is 0.1304 between folds, and the Bayesian MAD is 0.0843, a difference that is statistically significant by a simple sign test at a *p*-value too small for machine precision (easily *p* < 10^−100^).

### Entropy filtering improves reproducibility of both Pearson and Bayesian correlations

As described in the Introduction, the classical Relevance Networks algorithm begins by filtering out entities whose expression demonstrates low entropy. The purpose of this step is to avoid correlations that arise from a single sample or small set of “outliers.” Whether or not such an approach is appropriate is situation dependent. For example, if a subset of miRNAs were highly expressed only in glioblastoma multiforme tumours, and no others, such miRNAs would appear to have low entropy. (Remember, just five out of our 10,999 samples are for that disease.) We may not want to naively dismiss correlations among such miRNAs, as they arise from a clear disease relevance. Nevertheless, in the worst case, individual samples may be faulty and can create spurious correlations.

To test the effect of entropy filtering on both the Pearson and Bayesian correlations, we first computed the entropy of each miRNA’s expression (Fig 3A). In the original paper [12], it was suggested to discard the 5% of entities with lowest entropy (dashed red line). However, the appearance of the empirical entropy distribution suggested to us cut off around 10% (solid red line) would better separate entities with a “normal range” of entropies from those that appear unusually low. Hence, we chose 10% as our cut off, and defined miRNAs with entropies below that to be “low entropy” and the remainder to be “high entropy.” Fig 3B examines the relationship between miRNA expression and entropy. For the most part, the low entropy miRNAs also have very low expression. However, a small number of miRNAs with above average expression also have low entropy. The miRNA with the highest average expression that is still classified as low entropy is miR-205-3p, a miRNA with some known associations with cancer [32–34]. This miRNA is exceptionally high in two patient samples, one thymoma and one head or neck squamous cell carcinoma, where its expression levels of over 10,000 RPM are more than 100 times greater than in any other sample.

**Fig 3.**
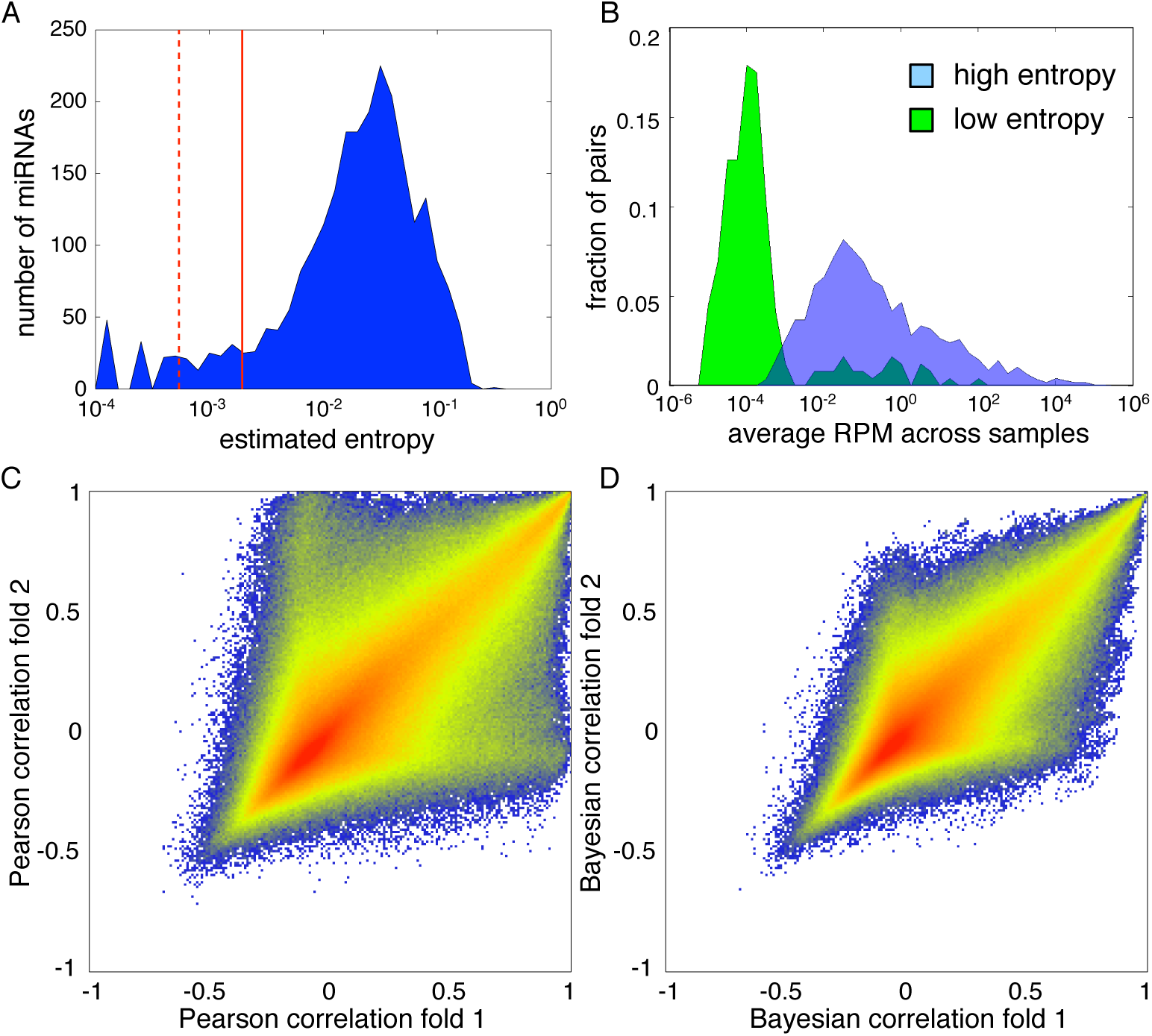
The effects of entropy filtering on Pearson and Bayesian correlations. (A) Empirical distribution of entropies of miRNAs’ expression across samples. Dashed red line indicates 5*^th^* percentile and solid red line indicates 10*^th^* percentile. (B) Empirical distribution of expression levels (average RPM across samples) for low entropy and high entropy miRNAs. (C) Comparison of Pearson grouped correlations from two halves of the data, when restricting attention to the high entropy miRNAs. (D) Comparison of Bayesian grouped correlations from two halves of the data, when restricting attention to the high entropy miRNAs.

Restricting attention to the high-entropy genes, and we recomputed the density scatterplots of Pearson correlations from the two halves of our data (Fig 3C), we see that the lines of exceptional correlations along the x- and y-axis are gone. (Compare to Fig 2B.) However, the overall qualitative shape of the point cloud remains, as do numerous miRNA pairs that have near +1 correlation in one half of the data and near zero correlation in the other half. Fig 3D shows the Bayesian correlations of the high-entropy miRNAs from each half of the data. There is little apparent change compared to Fig 2C, which includes the low entropy miRNAs. Perhaps surprisingly, entropy filtering does not improve the mean absolute deviation between the two halves of the data. For Pearson correlations restricted to high-entropy miRNAs, the MAD is 0.1305 (versus 0.1304 for all miRNAs), and for Bayesian correlations the MAD is 0.0894 (versus 0.0843). Although filtering eliminates some spurious correlations, it also eliminates many (correctly) zero correlations between low- or non-expressed miRNAs, driving the average error up.

### Bayesian Relevance Networks have lower false discovery rates

As Bayesian correlations between miRNAs match better between data folds than do Pearson correlations, we predicted that Bayesian Relevance Networks built based on each half of the data would agree better than classical Relevance Networks would. To test this hypothesis, we performed permutation testing on each half of the data, estimating null distributions for both the Pearson correlations and the Bayesian correlations. The results are shown in Fig 4A,B for analyzing all miRNA pairs (solid lines) and for analyzing high-entropy miRNAs only (dashed lines). The blue curves indicate the observed distributions of correlations on each half of the data, while the red curves indicate the estimated null distributions. For both Pearson and Bayesian correlations, there appear to be stronger positive correlations than would be predicted based on the null hypothesis of no statistical association between miRNAs. The shapes of the distributions estimated from each half of the data are in close agreement. There are more Pearson correlations at the highest levels (near 1) than there are Bayesian correlations—because of the tendency of the Bayesian approach to discount apparent correlations between low expression miRNAs. Filtering miRNAs based on entropy appears to have negligible impact on the distribution of Bayesian correlations, and a slightly larger, though still modest, effect on the Pearson correlations.

**Fig 4.**
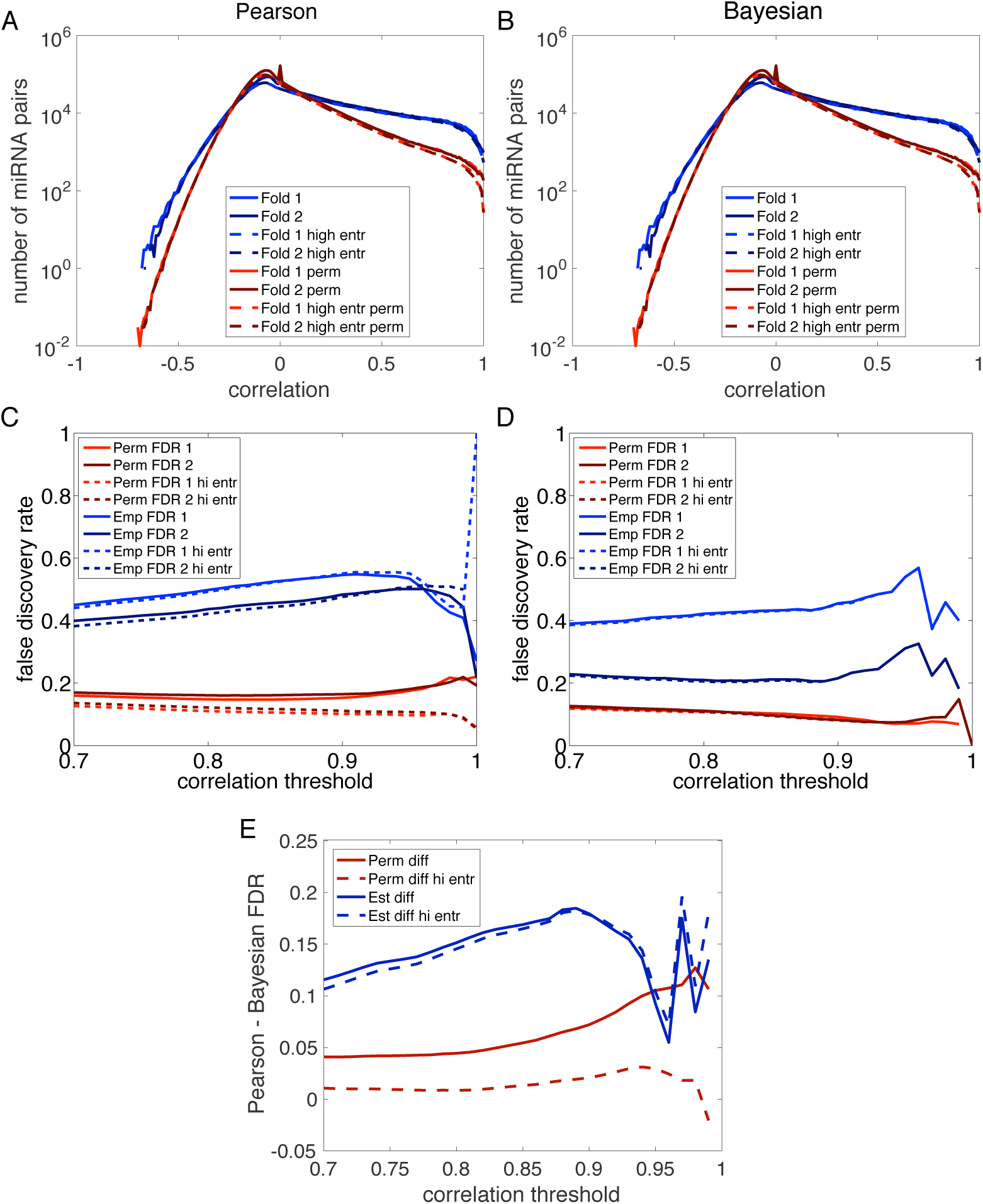
Permutation testing and agreement of Relevance Networks constructed based on Pearson or Bayesian correlations. (A) Empirical (blue) and permutation-based (red) distributions of Pearson correlations from each half of the data, comparing all pairs of miRNAs (solid lines) and comparing all pairs of high-entropy miRNAs (dashed lines). (B) Empirical (blue) and permutation-based (red) distributions of Bayesian correlations from each half of the data, comparing all pairs of miRNAs (solid lines) and comparing all pairs of high-entropy miRNAs (dashed lines). (C) Estimated false discovery rates (red, based on permutations) and empirical false discovery rates (blue, taking other half of the data as gold standard) at varying Pearson correlation thresholds. (D) Estimated false discovery rates (red, based on permutations) and empirical false discovery rates (blue, taking other half of the data as gold standard) at varying Bayesian correlation thresholds. (E) Difference between Pearson and Bayesian estimated and empirical false discovery rates.

Next, we constructed Relevance Networks at different correlation thresholds. At each threshold, we determined the number of miRNA pairs above threshold, as well as the expected number of such pairs under the null hypothesis. Based on these, we estimated the false discovery rate (FDR) for links in the Relevance Networks as a function of correlation threshold. At the same time, we compared the specific links constructed from each half of the data to the links in the other half. Links appearing in one half but not the other were labeled as putative false positives, and from these we constructed a second estimate of the FDR as a function of correlation threshold. The results are shown in Fig 4C,D,E and are radically different for Pearson and Bayesian approaches. Firstly, the Bayesian FDRs are almost uniformly better than the Pearson FDRs (Fig 4E); the only exception is at the threshold of *r* = 0.99, where the permutation-based estimate of Pearson FDR when restricting attention to high-entropy miRNAs is lower than the estimate for Bayesian FDR. When analyzing all miRNAs, the estimated Pearson FDRs from permutation testing hover around 0.2 for most correlation thresholds, whereas estimated Bayesian FDRs are smaller than 0.15. The empirical Pearson FDRs, based on comparing the networks obtained from each half of the data, are worse than 0.4 at all thresholds except *r* = 1, where there is substantial divergence between the estimates from the two data folds. The empirical Bayesian FDRs are somewhat different between the two folds of the data, but average to around 0.3 at most thresholds. The Bayesian FDR estimates either improve (drop) with increasing correlation threshold (permutation-based) or are relatively constant (based on data folds). This is a reasonable behaviour, as increasing the threshold intuitively means increasing stringency. Pearson FDRs sometimes decrease with increasing correlation threshold, but sometimes increase, depending on which estimate we consider and depending on the exact threshold level. When we restrict attention to the high-entropy miRNAs, we see that permutation-based estimates of FDR improve for the Pearson correlations. However, the empirical estimates of FDR based on comparing data folds do not improve. The FDRs of Bayesian Relevance Networks seem almost entirely immune to entropy filtering.

### A Bayesian Relevance Network describing co-expression of miRNAs across 10,999 patients with 33 types of cancer

Having established the soundness of the Bayesian Relevance Networks algorithm in the previous sections, we conclude the Results section by presenting the Bayesian Relevance Network obtained by analyzing the full dataset. We chose not to filter out miRNAs based on low entropy, so that we would not overlook potentially interesting connections, and because our results above suggest there would be little benefit. Accordingly, we computed all pairwise Bayesian correlations, and we performed 100 permutation computations to assess statistical significance. The empirical distributions of actual and permuted Bayesian correlations are shown in Fig 5A. As expected, we see many miRNA pairs that are highly correlated. However, high correlation can also be obtained by chance, as shown by the permutation testing. Even at a threshold of *r* = 0.99, which links just 61 miRNA pairs, our permutation testing suggests that 3.33 of those would be false positives.

**Fig 5.**
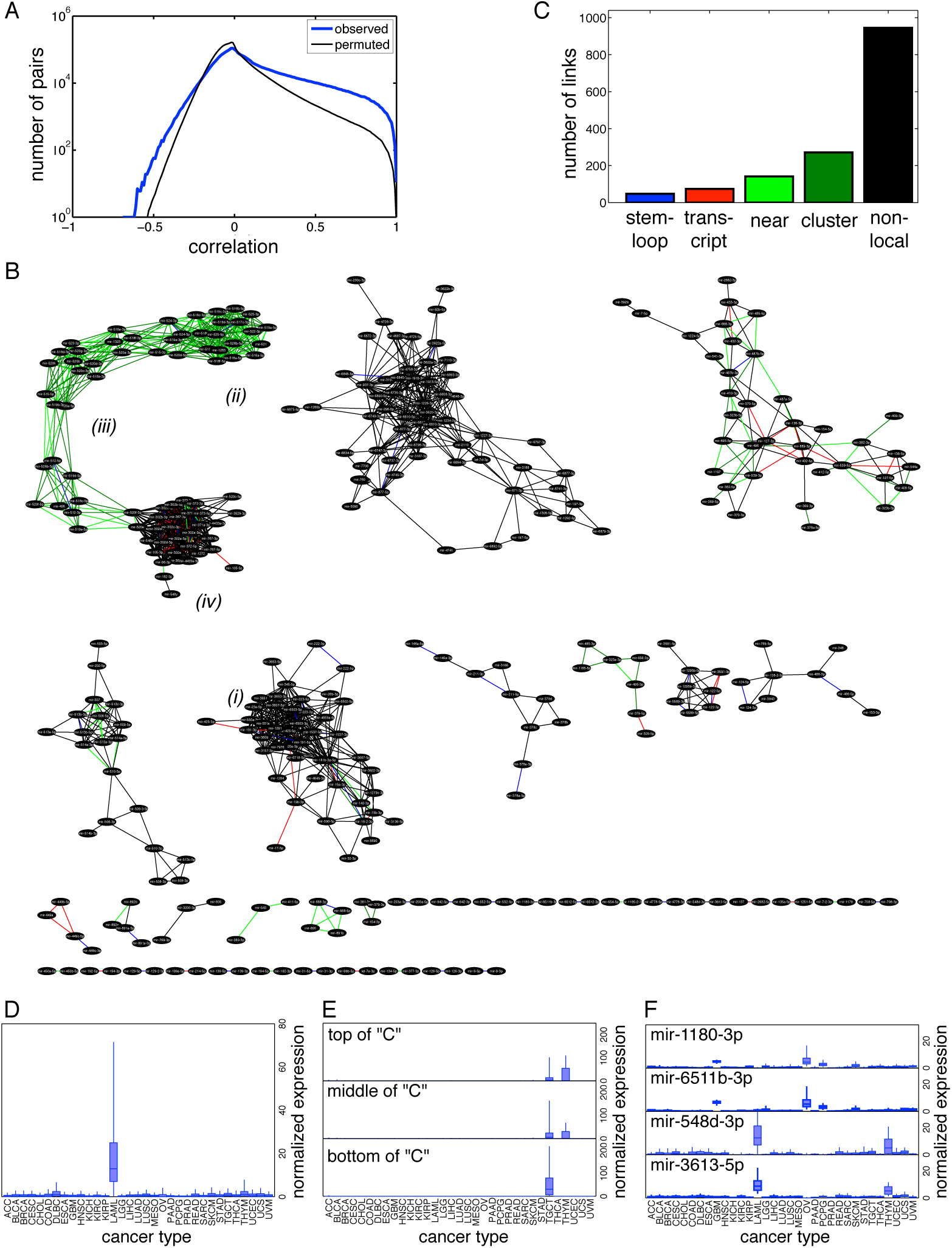
A Bayesian Relevance Network describing cross-cancer correlations between miRNAs. (A) Empirical distributions of Bayesian and permuted correlations. (B) The network obtained at a correlation threshold of *r* = 0.96. (C) Numbers of links with different locality relationships. (D) Normalized expression of miRNAs in the mostly-black subnetwork (*i*) near the center of the diagram in panel B. (E) Normalized expression of miRNAs at the top (*ii*), middle (*iii*), and bottom (*iv*) of the “C”-shaped subnetwork in the top left of panel A. (F) Normalized expression of four miRNAs participating in the only two non-local miRNA pairs in the relevance network.

We decided to construct the relevance network at the threshold *r* = 0.96. This gave us 1519 links between 342 distinct miRNAs, with an estimated 99.67 false positive links, or an empirical false discovery rate of 6.56%. We chose this level because it produced a large enough relevance network to see some interesting results, without letting the FDR grow too far out of control. The network is depicted in Fig 5B. We used Cytoscape [35] to construct the layout of the network. Links are colored by their locality: blue for miRNAs in the same pre-miRNA stem-loop, red for miRNAs in the same transcript, light green for miRNAs nearby on the genome, dark green for miRNAs in the same genomic cluster, and black for those not having any of those locality properties. As is typical for relevance networks, and indeed many types of biological networks, we observe connected components of widely varying sizes. Several major components have tens of miRNAs each and are heavily cross-connected, while there are also many isolated pairs of miRNAs connected by a single link. The majority of the links do not represent any locality relationship (Fig 5C).

A typical cluster is indicated by (*i*) in Fig 5B. Only a few links are related to genomic locale; most of the miRNAs are spread throughout the genome. miRNAs in this subnetwork are highly expressed in acute myeloid leukemia (TCGA code LAML) (Fig 5D). We found that many of the other connected subnetworks are also highly expressed in just one or a few cancer (or tissue) types.

A notable subnetwork is the “C”-shaped one in the upper left of the layout. This includes many miRNAs that are nearby on the genome (within 10kb) or at least within the same genomic cluster. However, the most densely connected part of the subnetwork, towards the bottom of the “C”, contains a mixture of stem-loop, transcript, local and non-local links. When we analyze miRNAs in three different parts of that network, we see different expression patterns (Fig 5E) The mostly-back cluster at the bottom is expressed almost exclusively in testicular germ cell tumors. At the opposite end of the “C”, the dense genomic cluster in green is expressed somewhat in testicular tumors but primarily in thymomas. miRNAs in between those two ends display a mixture of testicular tumor and thymoma expression. These miRNAs comprise the primate-specific C19MC miRNA cluster, which has normal functions in the placenta [36, 37]. This cluster’s roles in various cancers are still being worked out [38–41].

Although one must zoom in on the figure to see clearly, the vast majority of the links between isolated pairs of miRNAs do have some kind of locality relationship—unlike the majority of links in the network. Nearly half of the isolated miRNAs pairs are in the same stem-loop (11 of 23), five are in the same transcript, and five are nearby on the genome. Only two links are non-local, between miR-1180-3p and miR-6511b-3p, and between miR-548d-3p and miR-3613-5p (Fig 5F). These pairs show some evidence of cancer/tissue-specificity, with the first pair largely expressed in glioblastoma multiforme and ovarian cancer samples, and the latter pair largely expressed in acute myeloid leukemia samples and thymomas.

As a point of comparison, we computed a classical relevance network by computing mean expression levels for each miRNA within each cancer type in units of RPM, and then computing Pearson correlations between all pairs of miRNAs across the cancer types. One hundred permutation tests suggested that the minimum false discovery rate we could expect at any correlation threshold was over 15%, so we could not achieve the same error rate as in the Bayesian Relevance Network. Instead, we decided to compare the Bayesian and classical approaches when equalized to the same number of links. At a Pearson threshold of 0.9910775, the resulting Relevance Network had the same number of links (1519) as the Bayesian network, linking 308 distinct miRNAs. Many of these miRNAs and links are also present in the Bayesian Relevance Network, but many are not. Figure 6A shows a network depicting the difference between the Bayesian and Classical Relevance Networks. Black nodes and links are present in both networks. Green nodes and links are present only in the Bayesian network, and red nodes and links are present only in the classical network. The “C” shaped structure is present in both networks, with the classical network assigning additional miRNAs to the cluster at the bottom of the “C”. As we saw before, expression of these miRNAs is enriched in testicular germ cell tumors (Fig. 6B). The nodes added in the classical relevance network, however, have much lower expression levels. (The median expression level is zero reads.) The Bayesian approach does not “trust” their correlations enough to report them, but they may be legimate, and might have been included if greater sequencing depth revealed their expression levels more clearly. In general, miRNAs that were unique to the Classical Relevance Network had lower expression values than those uniquely in the Bayesian network, or those shared by both networks (Fig. 6C). In other parts of the difference network, we can see other subnetworks that were also present in the Bayesian Relevance Network, but which are now augmented by a few additional red nodes or links. But the Pearson analysis also failed to find many links reported by the Bayesian approach, as seen by the green nodes and links. Towards the bottom of the chart, we see a number of smaller subnetworks, most of which are unique to the Bayesian or Classical Relevance Networks. We were initially surprised that so many of the isolated pairs of linked miRNAs reported in the Bayesian Relevance Network (green doublets towards the bottom of Fig. 6A) are not present in the Pearson-based network. This is not because those links have low Pearson correlation estimates. Indeed, their Pearson correlations are uniformly larger than the Bayesian correlations. However, they were not high enough to reach the 0.9910775 threshold we needed so that the Classical Relevance Network would have as many links as the Bayesian network. In essence, other miRNA pairs with lower Bayesian correlations “leapfrogged” to even higher Pearson correlations, and thus were included in the Classical network.

**Fig 6.**
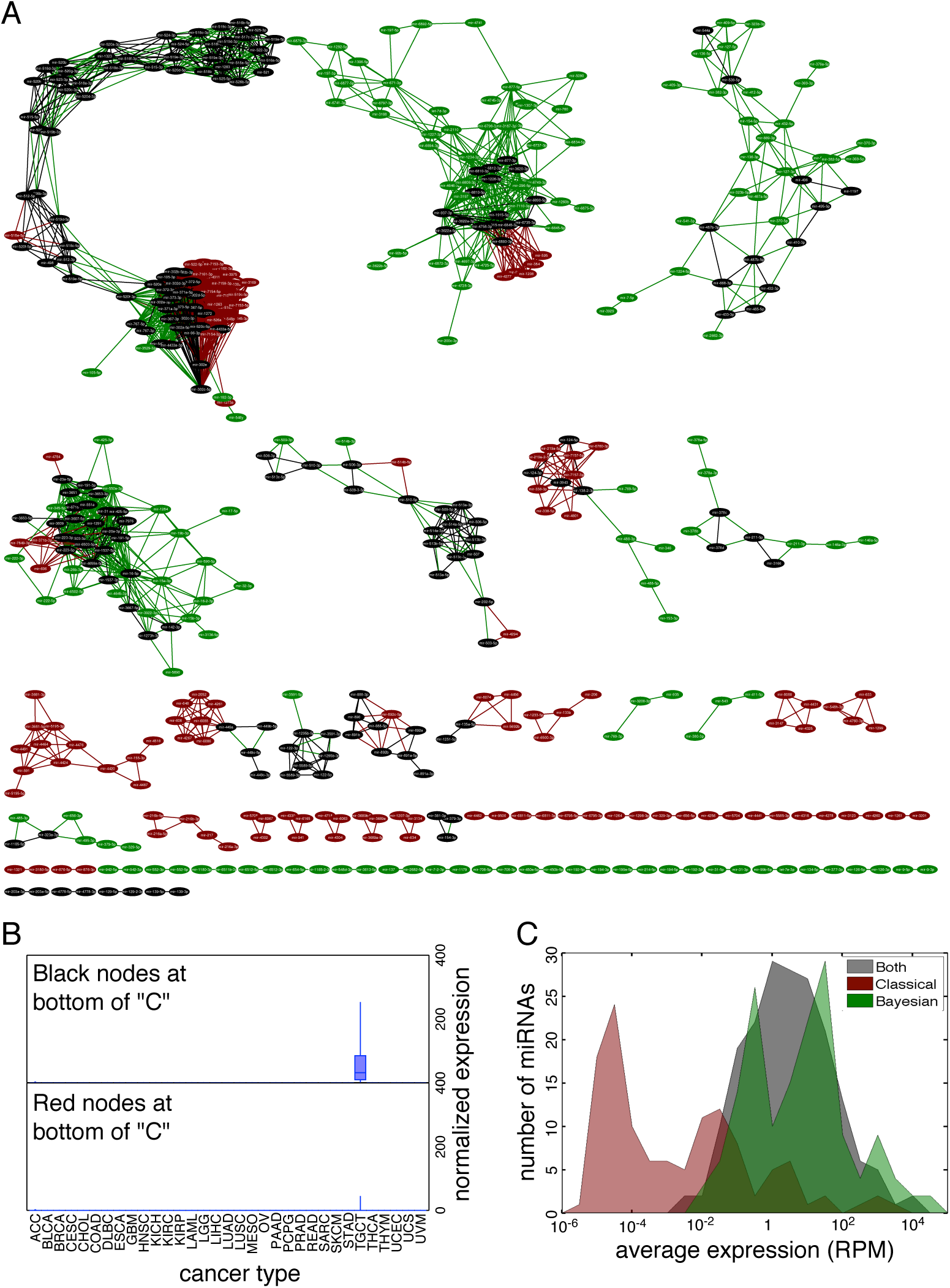
Difference between Bayesian and Classical Relevance Networks on cross-cancer co-expression. (A) Difference network in which black nodes and links are shared by both Bayesian and Classical Relevance Networks, red nodes and links are specific to the Classical network, and green nodes and links are specific to the Bayesian network. (B) Cancer-specific expression of shared and Classical-specific nodes at the bottom of the “C”-shaped structure. (C) Histogram of expression of shared, Bayesian-specific and Classical-specific miRNAs.

## Discussion

In this work, we have proposed Bayesian Relevance Networks as an update to the classical and widely-used Relevance Networks algorithm [12], with the aim of making it better suited to high-throughput sequencing data. Our approach accounts for the fact that sequence-based expression measurements can have widely varying precision, both for different entities (e.g., genes or miRNAs) and for different samples. It builds on our recent proposal for Bayesian correlation analysis [28], adding two main ingredients helpful for the construction of co-expression networks: 1) a method for estimating uncertainties in the expression levels in groups of samples; and 2) a permutation-testing scheme to assess statistical significance of Bayesian correlations. In testing on a large-scale miRNA expression dataset from The Cancer Genome Atlas [29], we found that Bayesian estimates of co-expression were more reproducible than the Pearson estimates used in the classical algorithm. As a consequence, we found that Bayesian Relevance Networks had lower false discovery rates than standard Relevance Networks. We also found that the entropy filtering step, with its additional and arbitrary cut off parameter, is unnecessary in the Bayesian approach, leading to a simpler algorithm over all. Although we focused on this single, large-scale dataset for demonstration and empirical evaluation, an important direction for future work is testing on other datasets. We suspect that one area where Bayesian Relevance Networks will be particularly helpful is in the analysis of single-cell RNA-seq data [42]. In such datasets, the average number of reads per gene are much smaller than for bulk RNA-seq data, and there can be great variability in the sequencing depths for each cell. This is exactly the situation where uncertainties in expression levels need to be considered, and where Bayesian approaches can provide a solution.

Bayesian Relevance Networks can be computed efficiently in both space and time, although there is a caveat regarding time efficiency. Regarding space, none of the computations are larger than *O*(*mn_tot_* + m^2^), where 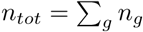 is the total number of samples. *O*(*mn_tot_*) is the size of the input count data, and *O*(*m*^2^) is the size of the output correlation matrix, so this performance is as good as possible. Regarding time, the complexity is *O*(*m^2^n_tot_*), because for all *O*(*m*^2^) pairs of entities, we need to perform various *O*(*n_tot_*) summations over samples or groups. For the miRNA dataset, in 10 repetitions on a 2013 MacBook Pro with 2.6 GHz i7 processor and 16 GB RAM, our Matlab implementation of the full Bayesian computation of all ≈ 3 million pairwise correlations took on average 205.2 ± 4.4 seconds. However, the vast bulk of that time was spent computing the numerator term *E_g_*(cov*_u_*(*p_ig_*,*p_j_*_g_)). Recall, this term represents the covariance, under our posterior belief, of the true expression levels of entities *i* and *j*. Because it involves a product of small fractions, this term is usually several orders of magnitude smaller than the other numerator term. Further, the exact same term appears in the permutation tests. When we omitted the second numerator term from all calculations, the computation sped up to 0.99 ± 0.02 seconds, and the results were virtually identical. This speed is comparable to that of our grouped Pearson calculations, which took 0.42 ± 0.01 seconds. Because of the large speed difference with and without the second numerator term, our Matlab and R codes for Bayesian correlations and permutations have an option allowing the user to skip that term. This is the time for computing all pairwise correlations. The permutation testing takes time approximately proportional to that multiplied by the number of repetitions. For 100 repetitions, for example, and skipping the second numerator term in the Bayesian correlations, the entire analysis can be completed in a matter of minutes.

In the broader context of co-expression network modeling, we view our primary contribution as emphasizing the importance of accounting for measurement uncertainty, and describing how that can be done in this era of sequencing-based expression measurement. As mentioned in the Introduction, Hughes *et al*. [1] paid significant attention to gene-specific measurement uncertainty in microarrays, using that information to discount correlations between genes. Much subsequent work on co-expression networks ommitted this issue, although some algorithms contain mechanisms that can be related to measurement uncertainty. For instance, the mutual information estimation in the ARACNE algorithm [18, 24] depends on a bivariate Gaussian kernel density estimate, with kernels placed on each observed pair of expression values. These kernels could be interpreted as representing a measurement uncertainty for each observation—although the authors do not talk about it in that manner. Another contribution of Hughes et *al*. [1] was to study the natural variability in expression of different genes, where they found that some genes are much more variable than others—as found in numerous other studies as well (e.g. [43, 44]). Our present algorithm accounts for only generic differences in entities and samples that arise because of differences in sequencing depth. However, given appropriate prior data, gene-specific expectations of expression levels or expression variability could be incorporated into our scheme through the Bayesian priors. Determining the best way to do this, and evaluating its impact on co-expression estimates, is an important topic for future work.

In our work, we have focused on incorporating uncertainty into the very simple, yet widely used, Pearson correlation metric. Much work on co-expression networks has explored other metrics for measuring similarity, such as the mutual information measures of Mutual Information Relevance Networks and the ARACNE algorithm [17, 18, 24] or the measures used in the WGCNA algorithm [22], which include weighted (i.e., exponentiated) versions of Pearson correlation, Spearman correlation and biweight midcorrelation. Our correlation metric could be immediately incorporated into WGCNA as an alternate fourth correlation measure. It could also be relatively easily incorporated into the ARACNE algorithm by observation-specific manipulation of kernel density bandwidths. An important direction for future work is to determine if accounting for measurement uncertainty increases the accuracy and reproducibility of algorithms such as these, as we found it to do for the Relevance Networks algorithm.

Co-expression sometimes suggests regulatory mechanisms, and so co-expression networks have been employed for the purpose of regulatory network estimation. This, however, brings up two related issues—direction of influence, and multiplicity of influence. While co-expression network construction is typically efficient, for those willing to pay the computational price, directed models such as Bayesian networks have been shown to be more accurate in some circumstances [45]. These models allow each gene to be regulated by multiple regulators, and, as generative models, can be used to make predictions about the outcome of perturbation experiments, for example. Static Bayesian networks have some limitations that co-expression networks do not, such as not permitting feedback loops—which are rife in biology in general and molecular networks in particular—due to the necessity of acyclic influence structure. But dynamic Bayesian networks can include feedbacks [46]. A final avenue for future research would be accounting for measurement noise in such a graphical model setting.

## References

1. Hughes TR, Marton MJ, Jones AR, Roberts CJ, Stoughton R, Armour CD, et al. Functional discovery via a compendium of expression profiles. Cell. 2000; 102: 109–126.

2. Stuart JM, Segal E, Koller D, Kim SK. A gene-coexpression network for global discovery of conserved genetic modules. Science. 2003; 302: 249–255.

3. Lee HK, Hsu AK, Sajdak J, Qin J, Pavlidis P. Coexpression analysis of human genes across many microarray data sets. Genome Res. 2004; 14: 1085–1094.

4. Baskerville S, Bartel DP. Microarray profiling of microRNAs reveals frequent coexpression with neighboring miRNAs and host genes. RNA. 2005; 11: 241–247.

5. Allocco DJ, Kohane IS, Butte AJ. Quantifying the relationship between co-expression, co-regulation and gene function. BMC Bioinformatics. 2004; 5: 1.

6. Michalak P. Coexpression, coregulation, and cofunctionality of neighboring genes in eukaryotic genomes. Genomics. 2008; 91: 243–248.

7. Lupien M, Eeckhoute J, Meyer CA, Wang Q, Zhang Y, Li W, et al. FoxA1 translates epigenetic signatures into enhancer-driven lineage-specific transcription. Cell. 2008; 132: 958–970.

8. Ponomarev I, Wang S, Zhang L, Harris RA, Mayfield RD. Gene coexpression networks in human brain identify epigenetic modifications in alcohol dependence. J Neurosci. 2012; 32: 1884–1897.

9. Chiou SH, Wang ML, Chou YT, Chen CJ, Hong CF, Hsieh WJ, et al. Coexpression of Oct4 and Nanog Enhances Malignancy in Lung Adenocarcinoma by Inducing Cancer Stem Cell–Like Properties and Epithelial–Mesenchymal Transdifferentiation. Cancer Res. 2010; 70: 10433–10444.

10. Hu S, Xu-Monette ZY, Tzankov A, Green T, Wu L, Balasubramanyam A, et al. MYC/BCL2 protein coexpression contributes to the inferior survival of activated B-cell subtype of diffuse large B-cell lymphoma and demonstrates high-risk gene expression signatures: a report from The International DLBCL Rituximab-CHOP Consortium Program. Blood. 2013; 121: 4021–4031.

11. Kubota Y, Shigematsu N, Karube F, Sekigawa A, Kato S, Yamaguchi N, et al. Selective coexpression of multiple chemical markers defines discrete populations of neocortical GABAergic neurons. Cereb Cortex. 2011; 21: 1803–1817.

12. Butte AJ, Tamayo P, Slonim D, Golub TR, Kohane IS. Discovering functional relationships between RNA expression and chemotherapeutic susceptibility using relevance networks. Proc Natl Acad Sci U S A. 2000; 97: 12182–12186.

13. Lennon NJ, Kho A, Bacskai BJ, Perlmutter SL, Hyman BT, Brown RH. Dysferlin interacts with annexins A1 and A2 and mediates sarcolemmal wound-healing. J Biol Chem. 2003; 278: 50466–50473.

14. Gomes LI, Esteves GH, Carvalho AF, Cristo EB, Hirata R, Martins WK, et al. Expression profile of malignant and nonmalignant lesions of esophagus and stomach: differential activity of functional modules related to inflammation and lipid metabolism. Cancer Res. 2005; 65: 7127–7136.

15. Elo LL, Järvenpää H, Orešič M, Lahesmaa R, Aittokallio T. Systematic construction of gene coexpression networks with applications to human T helper cell differentiation process. Bioinformatics. 2007; 23: 2096–2103.

16. Jiang W, Li X, Rao S, Wang L, Du L, Li C, et al. Constructing disease-specific gene networks using pair-wise relevance metric: application to colon cancer identifies interleukin 8, desmin and enolase 1 as the central elements. BMC Sys Biol. 2008; 2: 1.

17. Butte AJ, Kohane IS. Mutual information relevance networks: functional genomic clustering using pairwise entropy measurements. In: Pac Symp Biocomput. vol. 5; 2000. p. 26.

18. Basso K, Margolin AA, Stolovitzky G, Klein U, Dalla-Favera R, Califano A. Reverse engineering of regulatory networks in human B cells. Net Genet. 2005; 37: 382–390.

19. Schäfer J, Strimmer K, et al. A shrinkage approach to large-scale covariance matrix estimation and implications for functional genomics. Stat Appl Genet Mol Biol. 2005; 4: 32.

20. Faith JJ, Hayete B, Thaden JT, Mogno I, Wierzbowski J, Cottarel G, et al. Large-scale mapping and validation of Escherichia coli transcriptional regulation from a compendium of expression profiles. PLoS Biol. 2007; 5: e8.

21. Zhu D, Li Y, Li H. Multivariate correlation estimator for inferring functional relationships from replicated genome-wide data. Bioinformatics. 2007; 23: 2298–2305.

22. Langfelder P, Horvath S. WGCNA: an R package for weighted correlation network analysis. BMC Bioinformatics. 2008; 9: 559.

23. Acharya LR, Zhu D. Estimating an optimal correlation structure from replicated molecular profiling data using finite mixture models. In: Machine Learning and Applications, 2009. ICMLA’09. International Conference on. IEEE; 2009. p. 119–124.

24. Margolin AA, Nemenman I, Basso K, Wiggins C, Stolovitzky G, Dalla Favera R, et al. ARACNE: an algorithm for the reconstruction of gene regulatory networks in a mammalian cellular context. BMC Bioinformatics. 2006;7:S7.

25. AC’t Hoen P, Ariyurek Y, Thygesen HH, Vreugdenhil E, Vossen RH, de Menezes RX, et al. Deep sequencing-based expression analysis shows major advances in robustness, resolution and inter-lab portability over five microarray platforms. Nucleic Acids Res. 2008;36: e141–e141.

26. Git A, Dvinge H, Salmon-Divon M, Osborne M, Kutter C, Hadfield J, et al. Systematic comparison of microarray profiling, real-time PCR, and next-generation sequencing technologies for measuring differential microRNA expression. RNA. 2010; 16: 991–1006.

27. AC’t Hoen P, Friedländer MR, Almlof J, Sammeth M, Pulyakhina I, Anvar SY, et al. Reproducibility of high-throughput mRNA and small RNA sequencing across laboratories. Nat Biotechnol. 2013; 31: 1015–1022.

28. Sánchez-Taltavull D, Ramachandran P, Lau N, Perkins TJ. Bayesian Correlation Analysis for Sequence Count Data. PLoS ONE. 2016; 11: e0163595.

29. Weinstein JN, Collisson EA, Mills GB, Shaw KRM, Ozenberger BA, Ellrott K, et al. The cancer genome atlas pan-cancer analysis project. Nat Genet. 2013; 45: 1113–1120.

30. Grossman RL, Heath AP, Ferretti V, Varmus HE, Lowy DR, Kibbe WA, et al. Toward a shared vision for cancer genomic data. N Engl J Med. 2016; 375: 1109–1112.

31. Griffiths-Jones S, Saini HK, van Dongen S, Enright AJ. miRBase: tools for microRNA genomics. Nucleic Acids Res. 2008;36(Suppl 1):D154–D158.

32. Dahmke IN, Backes C, Rudzitis-Auth J, Laschke MW, Leidinger P, Menger MD, et al. Curcumin intake affects miRNA signature in murine melanoma with mmu-miR-205-5p most significantly altered. PLoS ONE. 2013; 8: e81122.

33. Jiang M, Zhang P, Hu G, Xiao Z, Xu F, Zhong T, et al. Relative expressions of miR-205-5p, miR-205-3p, and miR-21 in tissues and serum of non-small cell lung cancer patients. Mol Cell Biochem. 2013; 383: 67–75.

34. Vosgha H, Salajegheh A, Anthony Smith R, King-Yin Lam A. The important roles of miR-205 in normal physiology, cancers and as a potential therapeutic target. Curr Cancer Drug Targets. 2014; 14: 621–637.

35. Shannon P, Markiel A, Ozier O, Baliga NS, Wang JT, Ramage D, et al. Cytoscape: a software environment for integrated models of biomolecular interaction networks. Genome Res. 2003; 13: 2498–2504.

36. Tsai KW, Kao HW, Chen HC, Chen SJ, Lin WC. Epigenetic control of the expression of a primate-specific microRNA cluster in human cancer cells. Epigenetics. 2009; 4: 587–592.

37. Marie ND, Sayeda AA, Mohamed AK, Annick L, Philippe C, Gudrun ME, et al. The primate-specific microRNA gene cluster (C19MC) is imprinted in the placenta. Hum Mol Genet. 2010; p. ddq272.

38. Rippe V, Dittberner L, Lorenz VN, Drieschner N, Nimzyk R, Sendt W, et al. The two stem cell microRNA gene clusters C19MC and miR-371-3 are activated by specific chromosomal rearrangements in a subgroup of thyroid adenomas. PLoS ONE. 2010; 5: e9485.

39. Suzuki H, Takatsuka S, Akashi H, Yamamoto E, Nojima M, Maruyama R, et al. Genome-wide profiling of chromatin signatures reveals epigenetic regulation of MicroRNA genes in colorectal cancer. Cancer Res. 2011; 71: 5646–5658.

40. Augello C, Vaira V, Caruso L, Destro A, Maggioni M, Park YN, et al. MicroRNA profiling of hepatocarcinogenesis identifies C19MC cluster as a novel prognostic biomarker in hepatocellular carcinoma. Liver Int. 2012; 32: 772–782.

41. Kleinman CL, Gerges N, Papillon-Cavanagh S, Sin-Chan P, Pramatarova A, Quang DAK, et al. Fusion of TTYH1 with the C19MC microRNA cluster drives expression of a brain-specific DNMT3B isoform in the embryonal brain tumor ETMR. Nat Genet. 2014; 46: 39–44.

42. Jaitin DA, Kenigsberg E, Keren-Shaul H, Elefant N, Paul F, Zaretsky I, et al. Massively parallel single-cell RNA-seq for marker-free decomposition of tissues into cell types. Science. 2014; 343: 776–779.

43. Volfson D, Marciniak J, Blake WJ, Ostroff N, Tsimring LS, Hasty J. Origins of extrinsic variability in eukaryotic gene expression. Nature. 2006; 439: 861–864.

44. Choi JK, Kim YJ. Intrinsic variability of gene expression encoded in nucleosome positioning sequences. Nat Genet. 2009; 41: 498–503.

45. Werhli AV, Grzegorczyk M, Husmeier D. Comparative evaluation of reverse engineering gene regulatory networks with relevance networks, graphical gaussian models and bayesian networks. Bioinformatics. 2006; 22: 2523–2531.

46. Murphy K, Mian S. Modelling Gene Expression Data using Dynamic Bayesian Networks. Technical Report, Computer Science Division, University of California, Berkeley; 1999.

